# Consistency and identifiability of the polymorphism-aware phylogenetic models

**DOI:** 10.1101/718320

**Authors:** Rui Borges, Carolin Kosiol

## Abstract

Polymorphism-aware phylogenetic models (PoMo) constitute an alternative approach for species tree estimation from genome-wide data. PoMo builds on the standard substitution models of DNA evolution but expands the classic alphabet of the four nucleotide bases to include polymorphic states. By doing so, PoMo accounts for ancestral and current intra-population variation, while also accommodating population-level processes ruling the substitution process (e.g. genetic drift, mutations, allelic selection). PoMo has shown to be a valuable tool in several phylogenetic applications but a proof of statistical consistency (and identifiability, a necessary condition for consistency) is lacking. Here, we prove that PoMo is identifiable and, using this result, we further show that the maximum *a posteriori* (MAP) tree estimator of PoMo is a consistent estimator of the species tree. We complement our theoretical results with a simulated data set mimicking the diversity observed in natural populations exhibiting incomplete lineage sorting. We implemented PoMo in a Bayesian framework and show that the MAP tree easily recovers the true tree for typical numbers of sites that are sampled in genome-wide analyses.

## 1 Introduction

Phylogenies help to answer a multitude of questions regarding the origin of species, the tempo of evolution, the origin of particular traits and the processes (either neutral or selective) of molecular evolution (e.g. see (Pagel, 1999)). Phylogenies are therefore central to the study of patterns and processes by which evolution happens. However, phylogenies can only serve these purposes if they are correctly estimated. Fortunately, mathematical phylogenetics provides criteria that help us to assess whether a given phylogeny estimator is statistically sound. Statistical consistency and identifiability are two examples of such criteria.

In phylogenetic theory, a phylogenetic reconstruction method 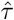 is statistically consistent under a model if it converges in probability to the true tree *τ* as the number of sites *S* of the sequence alignment increases indefinitely (Wald, 1949; Felsenstein, 1973), i.e.

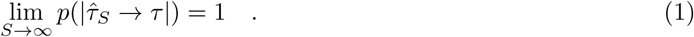

Identifiability is met if the model of evolution is uniquely characterized by the probability distribution it defines (Chang and Hartigan, 1991). An identifiable model is a necessary condition for consistency. More formal conditions for identifiability and consistency are described in Steel (1994), Chang (1996) and Steel (2013); these are revisited later in this article. Lack of statistical consistency has long been an aspect that phylogeneticists cared for. For example, some maximum parsimony methods were criticized for lacking statistical consistency early on (Felsenstein, 1978).

Simple phylogeny estimation methods using standard substitution model (e.g. Bayesian or maximum likelihood inference under the Jukes and Cantor (Jukes and Cantor, 1969), Kimura (Kimura, 1980) and Tajima and Nei (Tajima and Nei, 1984) substitution models) can be shown to enjoy statistical consistency (Chang, 1996). However, the same principle cannot be directly extended to more complex and general methods of tree inference that include rate variation across sites (Γ) and invariant sites (I). For such models of evolution, the model distribution is not fully understood, which complicates proving identifiability. Identifiability has been proven for the pure general time-reversible model (GTR, (Tavaré, 1986)). The GTR with rate variation (i.e. GTR+Γ) (Wu and Susko, 2010) also enjoys identifiability; however, a rigorous proof for the commonly utilized GTR+Γ+I is still lacking (Rogers, 2001; Allman et al., 2008; Chai and Housworth, 2011).

During the last years, alternative approaches to the classic nucleotide substitution models have been proposed. The polymorphism-aware phylogenetic models (PoMo) are such an example of an alternative approach for tree estimation that also accounts for incomplete lineage sorting (De Maio et al., 2013). While PoMo can be more broadly classified as a nucleotide substitution model; it adds a new layer of complexity by accounting for the population-level evolutionary processes (such as mutations, genetic drift, and selection) to describe the evolutionary process (De Maio et al., 2015; Schrempf et al., 2016; Borges et al., 2019). To do so, PoMo builds on a GTR-like mutation scheme and expands the {*A, C, G, T*} state-space to include polymorphic states, thereby accounting for current and ancestral intra-population variation. The latter aspect sets PoMo apart from the classic models of evolution, which traditionally only use a single representative DNA sequence per species.

PoMo has received substantial attention from the evolutionary community (see Mirarab et al. (2014), Szöllsi et al. (2015) and Leaché and Oaks (2017)). Several publications have employed it to solve a wide range of evolutionary questions, e.g., disentangling phylogenetic relationships among baboon species (Rogers et al., 2019), describing the phylogeographic history of great apes (Schrempf et al., 2016), estimating patterns of GC-bias and mutational biases (De Maio et al., 2013; Borges et al., 2019) and inferring the site-frequency spectrum (Schrempf and Hobolth, 2017; Borges et al., 2019) from population data.

All this raised the question of whether PoMo is a statistically consistent phylogeny estimator for phylogenetic data sets. Building upon the formal results provided by Steel (2013) and identifiability of the PoMo rate matrix and stationary distribution, we present a proof that the MAP topology (i.e. the tree topology that has the greatest posterior probability) is a statistically consistent estimate of the true tree. This result shows that PoMo is a statistically sound method of phylogenetic inference, and it provides validity for further investigations and uses of PoMo methods on real data sets.

**Table 1:** **Glossary**. The order of the symbols follows their first occurrence in the text.

| Symbol | Description |
| --- | --- |
| $\tau$ | tree topology |
| $\hat{\tau}$ | tree topology estimator |
| $S$ | number of sites |
| $a_i$ | allele $i$ |
| $\mathcal{A}$ | nucleotide base $\{A, C, G, T\}$ |
| $N$ | population size |
| $n$ | absolute frequency of an allele; relative frequency would be $n/N$ |
| $\mathbf{Q}$ | PoMo instantaneous rate matrix with elements $q$ |
| $\boldsymbol{\pi}$ | component of the mutation rate; stationary frequencies in the GTR model of evolution |
| $\boldsymbol{\rho}$ | component of the mutation rate; base exchangeabilities in the GTR model of evolution |
| $\boldsymbol{\sigma}$ | allelic selection coefficients |
| $k$ | normalization constant of the stationary vector |
| $\alpha$ | PoMo state |
| $K$ | number of alleles; $K = 4$ for PoMo |
| $L$ | number of taxa/sequences |
| $\mathbf{P}$ | transition matrix |
| $\chi$ | site pattern or, better, PoMo frequency patterns |
| $\eta$ | root vertex |
| $\epsilon = (\nu_1, \nu_2)$ | directed edge |
| $\boldsymbol{\omega}$ | branch lengths |
| $\boldsymbol{\theta}$ | set of PoMo parameters and branch lengths |

## 2 Polymorphism-aware phylogenetic models in a nutshell

PoMo assumes a Moran model (Moran, 1958) with *N* haploid individuals and defines the allele trajectory of a single locus with four possible alleles *a*_*i*_, where *i* ∈ 𝒜 = {*A, C, G, T*}. The evolution of this population in the course of time is described by a continuous time Markov chain with a discrete state-space defined by *N* and the four alleles. States are monomorphic (or boundary) if all the *N* individuals have the allele *i* {*Na*_*i*_}; or polymorphic, if two alleles are present in the population {*na*_*i*_, (*N − n*)*a*_*j*_} (Figure 1).

**Figure 1:**
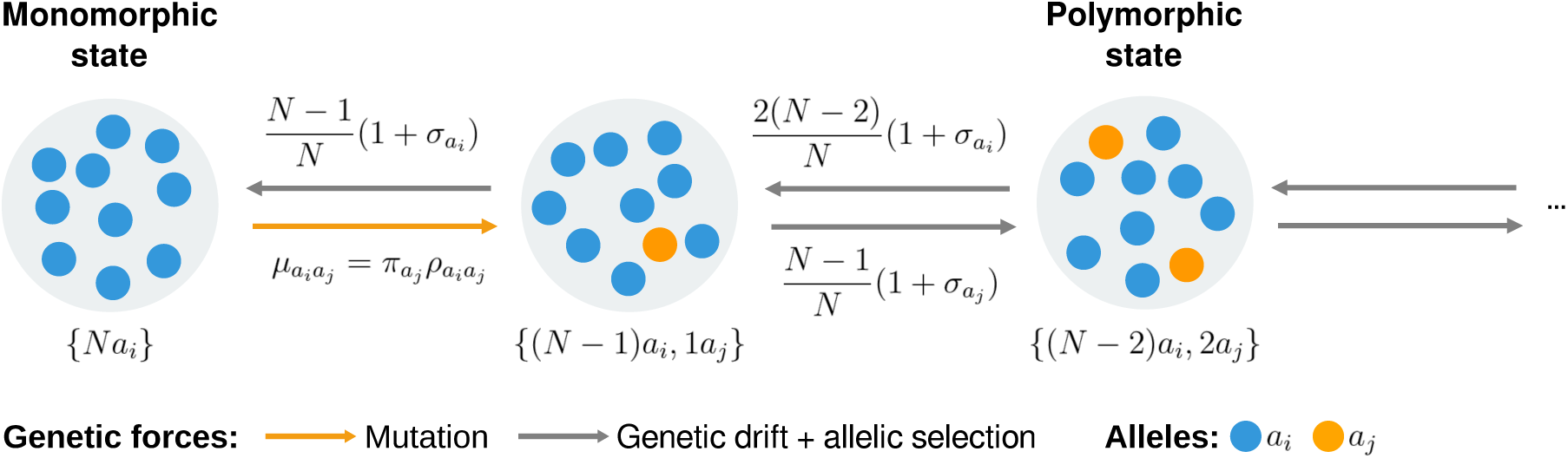
PoMo state-space and transition rates. The two alleles represent any of the four nucleotide bases A, C, G, and T. Orange and grey distinguish the role of mutation and genetic drift plus selection, respectively. The PoMo state-space includes monomorphic (or boundary states) {*Na*_*i*_} and polymorphic states {*na*_*i*_, (*N − n*)*a*_*j*_}. Monomorphic states interact with polymorphic states via mutation and polymorphic states reach monomorphic states via drift or selection. Between polymorphic states only drift and selection occur. PoMo is thus a particular case of the multivariate Moran model with boundary mutations and selection when four alleles (i.e. the four nucleotide bases) are considered.

A rate matrix ***Q*** describing the dynamic between the boundary and polymorphic states can be defined by considering several population-level processes. So far, PoMo includes mutation, genetic drift and allelic selection (Figure 1) (De Maio et al., 2015; Schrempf et al., 2016; Borges et al., 2019).

- Mutations are assumed to occur only in the boundary states {*Na*_*i*_} with mutation rates 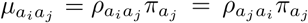. Mutation rates are thus modeled according to a GTR model of evolution (Tavaré, 1986), where ***π*** represents the stationary frequencies and ***ρ*** the six exchangeability terms. Similar interpretations of these parameters are still valid for the neutral PoMo, as ***π*** immediately informs the stationary frequencies of the monomorphic states. However, the stationary frequencies of the monomorphic states are no longer only defined by ***π*** in the more general PoMo with allelic selection (Borges et al., 2019) (Equation (3)). All in all, the GTR-like mutation scheme is a convenient strategy to obtain quantities of interest for PoMo.
- Genetic drift rules the allele frequency changes in a population. Genetic drift is modeled according to the Moran model, in which one individual is chosen to reproduce (i.e. to copy itself) and one to die in each time step (Moran, 1958). Therefore, the rates by which an allele with a starting frequency *n/N* is born or dies (i.e. the allele increases or decreases by one) are the same and equal to 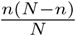. The allele frequency changes are thus neutral.
- Allelic selection may favor one allele over the other by differentiated reproductive success. The Moran machinery described previously can be adapted to model relative fitnesses by permitting that a given allele *a*_*i*_ has a fitness advantage/disadvantage 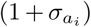 over the others (Durrett, 2008; Borges et al., 2019). ***σ*** refers to the vector of relative selection coefficients: i.e. a reference allele is chosen to have fitness 1, or selection coefficient 0.

Taking into account the population processes described so far, the PoMo instantaneous rate matrix ***Q*** is

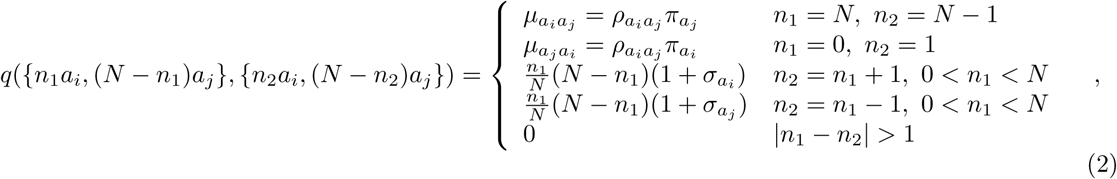

where *n*_1_ and *n*_2_ represent a shift in the allele frequencies. Frequency shifts larger than 1 are disallowed (last condition in equation (2)) making PoMo rate matrices typically sparse. The diagonal elements are defined such that the respective row sum is 0. The stationary distribution of PoMo is obtained by satisfying the condition ***ψQ*** = **0. *ψ*** is the normalized stationary vector and has the solution

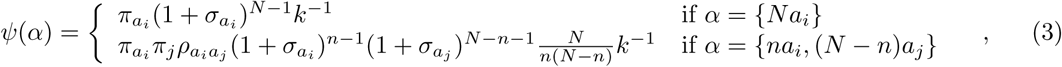

where *k* is the normalization constant and *α* a PoMo state (Borges et al., 2019).

When using PoMo to infer phylogenies, we make use of two important assumptions that are convenient for the proof of consistency presented here. Our first assumption is that sites evolve independently. Thus the probability that sequence *A* evolves to sequence *B* equals the product of the probability of evolutionary paths across all sites *S*. As a result, the site patterns created by the *L* species are independently and identically distributed (i.i.d.). The second assumption is that the evolutionary process is stationary and reversible with equilibrium measure ***ψ***. Proof of reversibility and stationarity have been provided by Schrempf et al. (2016) for the neutral PoMo model and by Borges et al. (2019) for the model with allelic selection.

## 3 Identifiability of PoMo

Identifiability is the inverse problem of finding the tree *τ* and the transition matrix ***P*** given just the probability of the various site patterns 𝓧 (or frequency patterns, in the case of PoMo) (Steel et al., 1998). Identifiability comes from the restrictions that must be placed on ***P*** for *τ* to be uniquely described by the probability of generating a pattern *χ*. These restrictions have been extensively studied (Chang and Hartigan, 1991; Steel, 1994; Chang, 1996; Steel et al., 1998).

Let us assume a tree *τ* with *η* representing the most recent common ancestor of *L* species (i.e. the root of the tree). The edges *∊* = (*v*_1_, *v*_2_) of *τ* are directed away from the root *η* in such a way that *v*_1_ lies between *η* and *v*_2_. If we attribute states to each vertex in the tree *τ*, beginning from the root *η* to all the descending vertices, we can represent the probability of generating pattern 𝓧 as

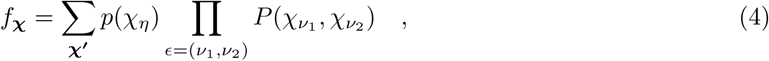

where 𝓧^**′**^ extends 𝓧 and represents the total assignment of states, and *p*(𝒳_*η*_) is the probability of state 𝒳_*η*_ at the root. The alphabet of PoMo has 4 + 6(*N -* 1) states and thus there are [4 + 6(*N -* 1)]^*L*^ possible frequency patterns 𝓧 in a tree with *L* leaves.

As PoMo makes some assumptions regarding the evolutionary process (Schrempf et al., 2016), we can further simplify equation (4): (i) frequency changes on edges are described by a continuous-time Markov process; (ii) the PoMo rate matrix ***Q*** is the same for all edges of the tree; and (iii) the distribution of frequency states at the root *p*(𝒳_*η*_) is simply the equilibrium distribution ***ψ***.

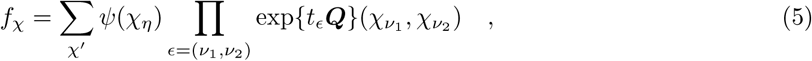

where *t*_*ϵ*_ is the length of edge *ϵ*. Conditions represented by (4) and (5) are also known as the Markov property, which is a necessary though not sufficient condition for identifiability.

Following Steel (1994); Steel et al. (1998), another condition for identifiability additional to (5) needs to hold: det(***P***) ≠ 0, 1, *-*1 for all edges *ϵ* in the tree and *ψ*(*α*) ≠ 0 for each PoMo state *α*. Then the tree *τ* can be uniquely recovered from the frequency patterns 𝓧 (Steel, 1994; Steel et al., 1998). By theorem 2.12 in Hall (2015), we can write that det(***P***) = exp{tr ***Q***} and therefore

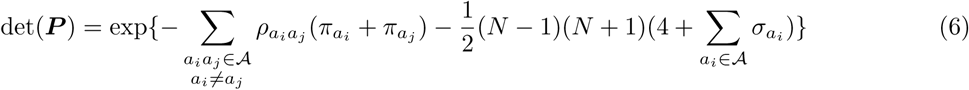

The selective pressures and the mutation rates, as modeled in PoMo, can only be real positive numbers. These rates cannot be 0 because they would represent very unlikely and biologically unreasonable scenarios: If 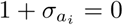 the individual carrying allele *a*_*i*_ would immediately die (i.e. an extremely deleterious allele); if 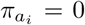 or 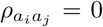 (both imply 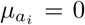) the allele *a*_*i*_ does not arise by mutation. In both situations, such allele should not be observed at all in the population, and one could simply use PoMo with *K -* 1 alleles, where *K* is the number of alleles in the population. Therefore, we can easily conclude that 0 *<* det(***P***_*ϵ*_) *<* 1 for all edges *ϵ* of *τ*.

Because we assume the equilibrium distribution of allele frequencies in the root, we need to check whether any elements of ***ψ*** can be equal to 0. The PoMo stationary distribution is defined for two different types of states *α*, the monomorphic (or fixed) and the polymorphic ones. As shown in equation (3), these states can only have 0 probability if any of the population parameters are 0. Therefore, we conclude that *ψ*(*α*) *>* 0 for all PoMo states *α*.

Summarizing, PoMo models respect the conditions for identifiability and we conclude that PoMo trees are uniquely identifiable by the frequency patterns they induce. Identifiability can be extended to the multivariate case by defining an alphabet 𝒜 over *K* alleles and a state-space of 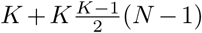 states. For simplicity we have considered the four-variate case, but the proofs shown here remain valid for the multivariate Moran model with allelic selection (Borges et al., 2019).

## 4 Consistency of the Bayesian PoMo

To evaluate the consistency property under PoMo, we have to prove that a given tree estimator converges in probability to the true tree as the number of sites *S* increases indefinitely. It has already been formally shown that the MAP tree (i.e. the tree topology that has the greatest posterior probability), estimated in a Bayesian framework, provides a statistically consistent estimator of the true tree (Steel, 2013). Consistency was proven under a wide variety of conditions (Steel, 2013), including tree inference from aligned sequences across the entire parameter range, and with the usage of general priors in models where the identifiability condition holds. Here, we show that these conditions are also met under PoMo.

Suppose we are given i.i.d. site patterns 𝓧 = (𝒳_1_, …, 𝒳_*S*_) generated by some unknown parameters (*τ*, ***θ***) and we wish to identify the topology *τ* from 𝓧 given prior densities on the set of fully resolved trees *T* and the continuous parameters ***θ***. These parameters include the branch lengths ***ω***, the mutations rates (defined by ***π*** and ***ρ***) and the selection coefficients ***σ***. Suppose we have a discrete prior probability distribution *p*(*τ*) on *T*, and, for each *τ ∈ T*, a continuous prior probability distribution on **Θ**(*τ*) with a probability density function *p*_*τ*_ (***θ***).

In particular, if the following four conditions hold:

- C1: *p*(*τ*) *>* 0;
- C2: the density *p*_*τ*_ (***θ***) is continuous, bounded and nonzero on **Θ**(*τ*);
- C3: the function ***θ*** → *p*_(*τ*,***θ***)_(𝓧) is continuous and nonzero on **Θ**(*τ*);
- C4: identifiability, guaranteeing that *τ* is uniquely identifiable by 𝓧.

Steel (2013) has shown that

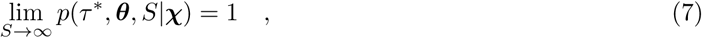

where *p*(*τ* ^*^, ***θ***, *S*|𝓧) is probability that the MAP correctly selects *τ* ^*^ for a i.i.d. sequence (𝒳_1_,, 𝒳_*S*_) generated by (*τ* ^*^, ***θ***). In other words, the MAP maximizes the posterior probability *p*(*τ*)*E*_***θ***_[*p*(𝓧|*τ*, ***θ***)] where *E*_***θ***_ is the expectation of the likelihood with respect to the prior probability of **Θ**(*τ*).

Consistency can then be inherently gained by Bayesian inference. PoMo can be easily placed in a Bayesian framework. Indeed, we can easily meet conditions 1 and 2 in standard Bayesian phylogenetic inference software (e.g. BEAST (Bouckaert et al., 2019) and RevBayes (Höhna et al., 2016)). C1 requires that a nonzero prior is set on the tree topology, which can be easily implemented by setting a uniform prior. If ***θ*** takes the usual exponential/gamma priors on the branch lengths and rate parameters (***ρ, σ*** and ***ω***) or the Dirichlet distribution on ***π*** (which can actually be generated from a set of *K*-independent gamma random variables), condition C2 is met. Conditions C3 and C4 hold for all Markov processes on any tree with pendant edges of positive lengths (i.e. on any binary metric-tree) for which identifiability was proven. Therefore, as shown in the previous section, C3 and C4 hold for PoMo. Consequently, the MAP tree under PoMo is a consistent estimator of the species tree.

## 5 A simulation-based example of consistency with PoMo

Consistency guarantees the identification of the correct parameter values with infinite sequence lengths. In real data situations, the sequence length is finite as is the running time. We have nevertheless tested the consistency of the tree topology for the Bayesian PoMo estimator using simulated population data sets.

We simulated alignments of 10 000 sites under PoMo using a phylogeny of five populations as shown in figure 2A with relative branch lengths defined in such a way that we have two closely and two distantly (i.e. twice the expected divergence) related populations. We simulated two scenarios I and II by mimicking fast and slowly evolving populations (expected divergence *D* equal to 0.3 and 0.02 substitutions per site, respectively) with expected heterozygosity *H*_*e*_ as observed in *Drosophila simulans* populations and among great apes (0.0015 and 0.018, respectively) (Begun et al., 2007; Prado-Martinez et al., 2013).

**Figure 2:**
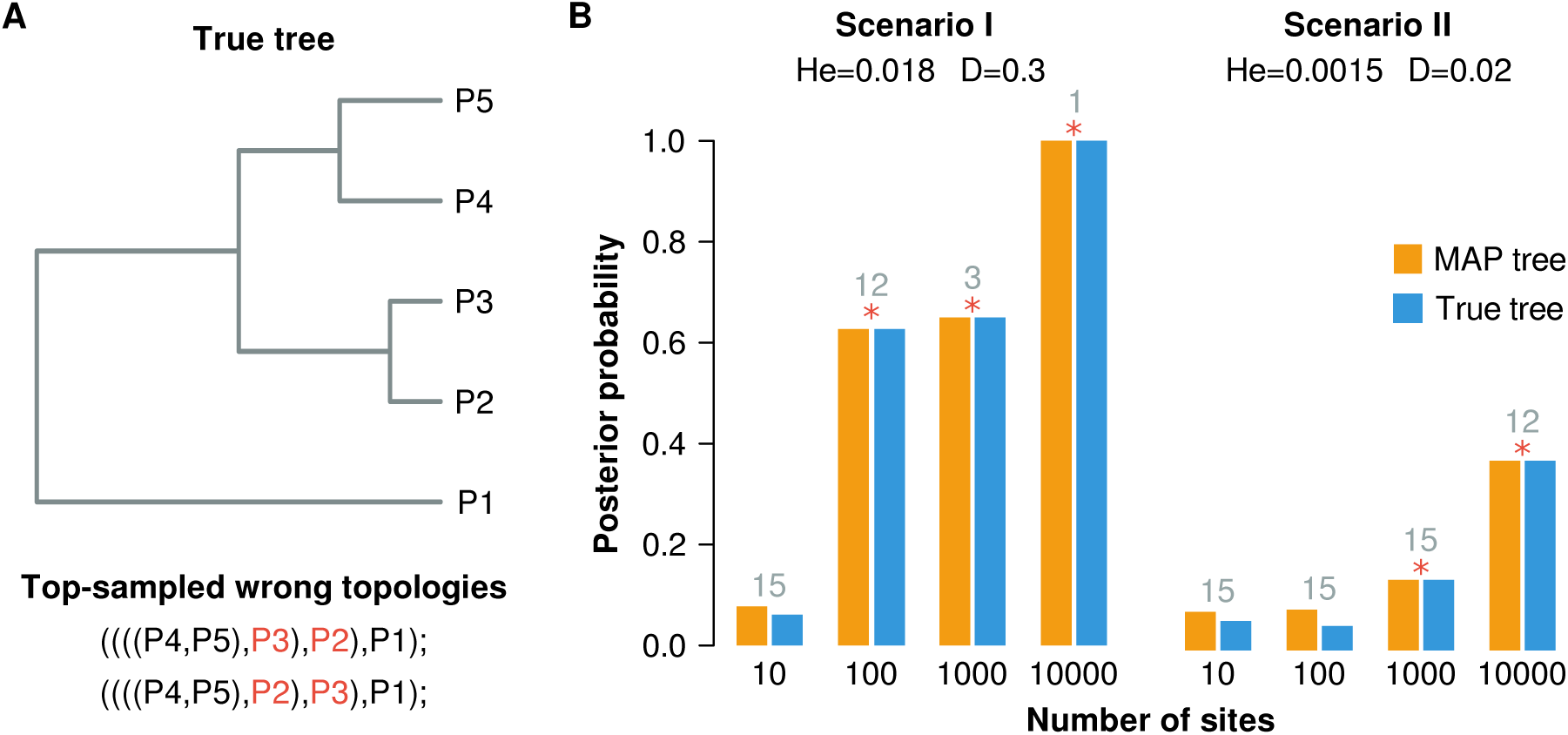
Consistency of the Bayesian PoMo. **(A)** Topology (and relative branch lengths) used for simulations and the two top-visited wrong topologies sampled from the posterior. These alternative topologies follow the expected tree discordance due to incomplete lineage sorting between populations 2 and 3. **(B)** Posterior probabilities of the MAP and true topologies for the two simulated scenarios I and II. The expected heterozygosity *H*_*e*_ and divergence *D* used to simulate each scenario were taken from natural populations of *D. simulans* (scenario I (Begun et al., 2007)) and great apes (scenario II (Prado-Martinez et al., 2013)). The asterisk indicates that the MAP tree corresponds to the true tree. Numbers in grey indicate the number of sampled posterior topologies.

To test for consistency, we created four alignments including the first 10, 100, 1000, 10 000 sites. We fed these alignments to RevBayes (Höhna et al., 2016) and performed standard Bayesian phylogeny estimation on them. We ran PoMo for two chains and 20 000 generations, keeping every 10th iteration. A burning period of 10% was defined by checking mixing, convergence, and auto-correlation of the MCMC chains.

We observed that MAP already recovers the true tree for the average gene length of 1000 sites (Figure 2B). These simulated scenarios complement our proofs that the MAP is a consistent estimator of the true tree. We observed further that populations with more heterozygous alignments (i.e. scenario I) converge to the true tree faster. The effect of incomplete lineage sorting is evident in these examples, as the top-sampled wrong topologies (Figure 2A) cluster the closely related populations 3 and 2 with the clade including population 4 and 5. These topologies are, for a fewer number of sites, the MAP trees of scenarios I and II.

## 6 Conclusions

Here we prove that PoMo is identifiable and further that the MAP is a statistically consistent estimator of the species tree when PoMo is placed in a Bayesian framework. This is the first time identifiability and consistency were shown for the polymorphism-aware phylogenetic models.

Identifiability of PoMo can be easily extended to important, more general models. Identifiability should be kept for generalizations that work on the standard PoMo instantaneous rate matrix: This is, for example, the case with balancing selection acting along with genetic drift. More complicated extensions of the PoMo models would include the joint inference of gene trees with the species trees; currently, PoMo directly estimates the species tree. Steel (2013) suggested that identifiability and consistency should still apply, as the gene trees can be viewed as nuisance parameters. A formal proof for this statement, especially for the case of gene trees undergoing gene gain, loss, and transfer, is missing.

The consistency property, as already mentioned, says nothing about the performance of the method in real contexts, where data is finite. However, consistency is a desirable property, especially for the type of data PoMo is applied to: population-scale and genome-wide. This data is costly and technically difficult to obtain and indeed the data sets available in the literature are few (e.g. humans (Altshuler et al., 2010), great apes (Prado-Martinez et al., 2013) fruit flies (Lack et al., 2015) and *Arabidopsis* (Long et al., 2013)). It is only natural to expect that higher samples sizes are repaid with less erroneous estimates of model parameters, including the species tree.

Our simulated examples, though simple, show that the MAP recovers the true tree even when the expected heterozygosity is very low and with incomplete lineage sorting is present. ILS is a very well-known cause of discordance between gene and species trees (Maddison and Knowles, 2006), by affecting the probability of visited wrong topologies. Statistical consistency is thus a desirable property for phylogeny estimation on closely related populations. As we have seen, the MAP tree corresponds already to the true topology for 1000 sites, even for less diverse species.

As future work, we would want to explore the sequence length requirement under PoMo. This is essentially the sequence length that a phylogeny reconstruction method needs to recover the true tree with a considerably small error (Atteson, 1999). A low sequence length requirement is a condition for a computationally efficient method. In particular, it would be important to determine whether PoMo is a fast converging method (i.e. a method that requires a sequence length of only *𝒪*(*poly*(*n*))).

## Acknowledgements

This work has been funded by the Vienna Science and Technology Fund (WWTF) through project MA16-061. We thank Bastien Boussau for helping with RevBayes and Lynette Mikula for useful comments and suggestions on an earlier version of this manuscript.

